# Prevalence and correlates of phenazine resistance in culturable bacteria from a dryland wheat field

**DOI:** 10.1101/2021.11.23.469799

**Authors:** Elena K. Perry, Dianne K. Newman

**Affiliations:** Division of Biology and Biological Engineering, California Institute of Technology, Pasadena, CA 91125, USA; Division of Geological and Planetary Sciences, California Institute of Technology, Pasadena, CA 91125, USA

## Abstract

Phenazines are a class of bacterially-produced redox-active natural antibiotics that have demonstrated potential as a sustainable alternative to traditional pesticides for the biocontrol of fungal crop diseases. However, the prevalence of bacterial resistance to agriculturally-relevant phenazines is poorly understood, limiting both the understanding of how these molecules might shape rhizosphere bacterial communities and the ability to perform risk assessment for off-target effects. Here, we describe profiles of susceptibility to the antifungal agent phenazine-1-carboxylic acid (PCA) across more than 100 bacterial strains isolated from a wheat field where PCA producers are indigenous and abundant. We find that Gram-positive bacteria are typically more sensitive to PCA than Gram-negative bacteria, but that there is also significant variability in susceptibility both within and across phyla. Phenazine-resistant strains are more likely to be isolated from the wheat rhizosphere, where PCA producers are also more abundant, compared to bulk soil. Furthermore, PCA toxicity is pH-dependent for most susceptible strains and broadly correlates with PCA reduction rates, suggesting that uptake and redox-cycling are important determinants of phenazine toxicity. Our results shed light on which classes of bacteria are most likely to be susceptible to phenazine toxicity in acidic or neutral soils. In addition, the taxonomic and phenotypic diversity of our strain collection represents a valuable resource for future studies on the role of natural antibiotics in shaping wheat rhizosphere communities.

**Importance:** Microbial communities contribute to crop health in important ways. For example, phenazine metabolites are a class of redox-active molecules made by diverse soil bacteria that underpin the biocontrol of wheat and other crops. Their physiological functions are nuanced: in some contexts they are toxic, in others, beneficial. While much is known about phenazine production and the effect of phenazines on producing strains, our ability to predict how phenazines might shape the composition of environmental microbial communities is poorly constrained; that phenazine prevalence in the rhizosphere is predicted to increase in arid soils as the climate changes provides an impetus for further study. As a step towards gaining a predictive understanding of phenazine-linked microbial ecology, we document the effects of phenazines on diverse bacteria that were co-isolated from a wheat rhizosphere and identify conditions and phenotypes that correlate with how a strain will respond to phenazines.

## Introduction

Diverse microorganisms produce natural antibiotics that can kill or inhibit the growth of other microbes (Bérdy, 2012; Granato *et al*., 2019). Several such compounds have been commercialized as antimicrobial drugs for the treatment of infections, beginning with penicillin in the 1940s (Aminov, 2010; Hutchings *et al*., 2019). Unfortunately, the selective pressures exerted by the widespread use of antibiotics in medicine and agriculture have led to worrisome increases in the prevalence of antimicrobial resistance among human and animal pathogens over the past several decades (Davies and Davies, 2010; Manyi-Loh *et al*., 2018). Yet while this repercussion of human antibiotic use has been well documented, comparatively little is known about the ecological effects of microbially-produced antibiotics in natural environments (Aminov, 2009; Sengupta *et al*., 2013). Recent studies have suggested that natural antibiotics may serve a variety of functions for their producers beyond the suppression of competing microbes (Davies, 2006; Davies *et al*., 2006), including nutrient acquisition (Wang and Newman, 2008; McRose and Newman, 2021), conservation of energy in the absence of oxygen (Glasser *et al*., 2014; Glasser *et al*., 2017), and cell-cell signaling (Dietrich *et al*., 2006; Linares *et al*., 2006). At the same time, toxicity to one or more microorganisms is by definition a feature of these molecules, but the extent to which this trait shapes their influence on microbial communities is unclear (Demain and Fang, 2000; Davies and Ryan, 2012; Bérdy, 2012). In addition, for many if not most natural antibiotics, the determinants and prevalence of susceptibility or resistance to their toxicity remain unknown or poorly characterized (Handelsman and Stabb, 1996). These gaps in knowledge hinder our ability to understand and predict the impacts of these metabolites on microbial communities of interest.

One environmental context in which natural antibiotics are thought to be particularly abundant and ecologically relevant is the rhizosphere—the narrow plane of soil immediately adjacent to plant roots (Haas and Défago, 2005; Mavrodi *et al*., 2012; Tyc *et al*., 2017). Natural antibiotics such as phenazines and 2,4-diacetylphloroglucinol have been directly detected in the rhizospheres of multiple crops, including wheat, potato, and sugar beet (Thomashow *et al*., 1990; Bergsma-Vlami *et al*., 2005; Mavrodi *et al*., 2012), and phenazines have been shown to increase the fitness of their producers when competing with other microbes in the rhizosphere (Mazzola *et al*., 1992; Yu *et al*., 2018). Production of these molecules has also been demonstrated to underpin the ability of certain bacteria to control fungal crop diseases (Thomashow and Weller, 1988; Thomashow *et al*., 1990; Haas and Keel, 2003; Mazurier *et al*., 2009; Yu *et al*., 2018), further indicating that natural antibiotics can act as agents of microbe-microbe antagonism in the rhizosphere. As a result of this activity, phenazine-producing *Pseudomonas* strains have received attention as potential biocontrol agents that could serve as a more sustainable alternative to traditional synthetic pesticides in agriculture (Handelsman and Stabb, 1996; Haas and Keel, 2003). However, several challenges remain with respect to the practical application of these strains, including inconsistent efficacy under field conditions (Haas and Keel, 2003; Haas and Défago, 2005), limited understanding of the mechanisms and evolutionary dynamics of resistance to phenazines (Mazzola *et al*., 1995; Handelsman and Stabb, 1996), and the possibility of off-target effects (Haas and Keel, 2003).

Importantly, with regard to the latter concern, phenazines are toxic not only to fungi, but also to some bacteria (Baron and Rowe, 1981; Turner and Messenger, 1986; Costa *et al*., 2015). Yet while the utility of phenazine-producing pseudomonads for biocontrol of fungal crop diseases has been extensively investigated, the potential impact of these strains on non-target bacterial residents of the rhizosphere is less well understood. One study found that inoculation of *Pseudomonas* biocontrol strains shifted the rhizosphere community of maize seedlings, pushing the ratio of Gram-positive to Gram-negative bacteria in favor of the latter; however, this analysis was based on profiling colony growth rates and whole-cell fatty acids from pooled cultured isolates, greatly limiting the taxonomic resolution and making it difficult to rule out whether the Gram-negative biocontrol strains might themselves have directly contributed to the shift (Kozdrój *et al*., 2004). On the other hand, at least three studies found no notable or consistent effects of introduced *Pseudomonas* species on the rhizosphere bacterial communities of wheat or potato (Gagliardi *et al*., 2001; Bankhead *et al*., 2004; Roquigny *et al*., 2018), albeit the studies in wheat employed methods with limited discriminatory power (namely, carbon source utilization profiling and terminal restriction fragment length polymorphism). Given these mixed results and the lack of fine-grained spatial or taxonomic resolution in most studies on this topic, whether phenazine-producing bacteria actively shape the surrounding rhizosphere bacterial community through antibiosis, potentially at the micron or millimeter scale, remains an open question.

In addition to the lack of clarity regarding the effects of phenazines on rhizosphere bacterial communities, the taxonomic and physiological correlates of phenazine resistance in bacteria remain incompletely understood. The toxicity of phenazines is generally ascribed to the generation of reactive oxygen species (ROS) and interference with respiration (Hassan and Fridovich, 1980; Baron and Rowe, 1981; Voggu *et al*., 2006; Perry and Newman, 2019). Previous studies have suggested that efflux pump expression, cell permeability, oxidative stress responses, and the composition of the respiratory electron transport chain can affect bacterial susceptibility to phenazines (Voggu *et al*., 2006; Khare and Tavazoie, 2015; Noto *et al*., 2017; Wolloscheck *et al*., 2018; Perry and Newman, 2019; Meirelles *et al*., 2021). In addition, a comparison of 14 bacterial strains indicated that Gram-negative bacteria as a group may be more resistant to phenazine toxicity than Gram-positive bacteria (Baron and Rowe, 1981). However, all of these studies focused on a specific phenazine, pyocyanin, that is particularly toxic (Meirelles and Newman, 2018) and best known for its role as a virulence factor during infections of humans and animals (Lau *et al*., 2004; Liu and Nizet, 2009). Whether the same observations hold true for more agriculturally-relevant phenazines such as phenazine-1-carboxylic acid (PCA) (Thomashow and Weller, 1988; Thomashow *et al*., 1990; Mavrodi *et al*., 2012; Dar *et al*., 2020) is unknown.

In this study, we set out to lay a foundation for addressing unresolved questions about the ecological impact of phenazine toxicity in the rhizosphere by determining the prevalence of phenazine resistance among bacteria isolated from a wheat field in the Inland Pacific Northwest, a region where phenazine production and the biocontrol potential of indigenous *Pseudomonas* species have been studied for decades (Thomashow and Weller, 1988; Thomashow *et al*., 1990; Mazzola *et al*., 1992; Mavrodi *et al*., 2012). We designed a culture-based assay to measure sensitivity to PCA, which is the best-studied and one of the most abundant phenazines in this environment (Thomashow *et al*., 1990; Mavrodi *et al*., 2012; Dar *et al*., 2020). We also performed full-length 16S rRNA gene sequencing of our isolates in order to assess the relationship between taxonomy and PCA resistance. Finally, to assess potential physiological correlates of PCA resistance, we measured PCA reduction rates for a subset of phenotypically-diverse strains and investigated the effects of broad-spectrum efflux pump inhibitors on growth in the presence of PCA.

## Results

### Taxonomic diversity of culturable bacteria from dryland wheat rhizospheres and bulk soil

A total of 166 strains of bacteria were isolated from 12 soil samples collected from a wheat field at Washington State University’s Lind Dryland Research Station in early August 2019, shortly after the wheat harvest. The samples comprised 4 replicates each of wheat rhizosphere (“Wheat”), bulk soil collected between planted rows (“Between”), and bulk soil from a virgin patch of uncultivated soil adjacent to the field (“Virgin”). Full-length 16S rRNA gene sequencing revealed that the isolates represented 24 genera across 4 phyla: Actinobacteria, Bacteroidetes, Firmicutes, and Proteobacteria. The vast majority of isolates from the bulk soil samples (both Between and Virgin) were Actinobacteria or Firmicutes. In Wheat samples, on the other hand, the combined proportions of these two phyla were lower, accounting for 18-50% and 10-25% of isolates respectively, while Proteobacteria accounted for approximately 25-60% of isolates depending on the replicate, and 1-3 strains of Bacteroidetes were also detected in each replicate (5-12% of isolates) (Fig. 1).

**Figure 1:**
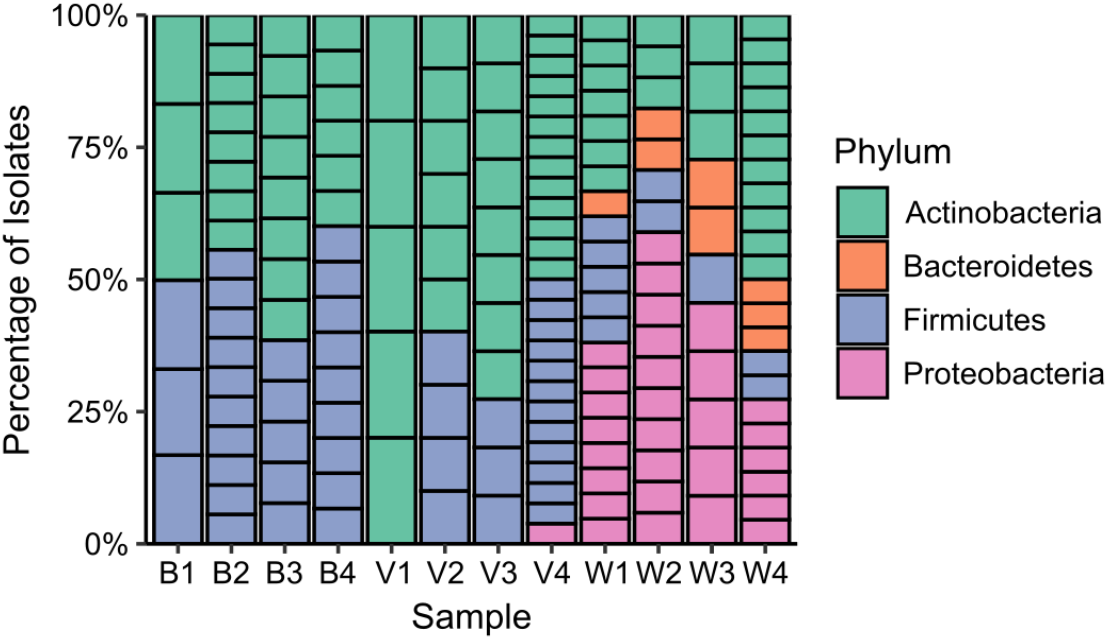
Taxonomic distribution of bacterial isolates from wheat rhizosphere and bulk soil samples. This plot depicts the proportion of isolates from each sample that belonged to the 4 represented phyla. Each column represents one soil sample, and each box within the columns represents an individual isolate colored by the phylum to which it belongs (e.g. 6 boxes comprising one column indicates that 6 strains were isolated from that sample). B = Between (bulk soil), V = Virgin (bulk soil), W = Wheat (rhizosphere).

### Taxonomic and spatial distribution of PCA resistance phenotypes

We screened our isolates for resistance to PCA at both circumneutral and acidic pH (7.3 and 5.1, respectively), because the toxicity of PCA to diverse organisms is known to vary depending on pH (Brisbane *et al*., 1987; Cezairliyan *et al*., 2013), and because the bulk soil and rhizosphere pH of wheat fields in the Inland Pacific Northwest can vary by >3 units (pH 4.3-8.0) depending on geographic location and fertilizer treatment status (Smiley and Cook, 1973). The pH-dependency of PCA toxicity has been attributed to the fact that the deprotonated form of PCA is negatively charged (Fig. S1); the negative charge on bacterial cell walls and the outer membrane of Gram-negative bacteria (or the negative membrane potential of eukaryotic cells) likely hinders uptake of this species. The protonated form of PCA, on the other hand, is neutral and presumably can passively diffuse across cell membranes given the small size and hydrophobic nature of the molecule (Price-Whelan *et al*., 2006; Cezairliyan *et al*., 2013). The pKa of PCA is 4.24 at 25 °C; thus, at pH 7, only 0.17% of PCA in solution is protonated compared to 14.8% at pH 5, typically leading to greater toxicity at the lower pH (Brisbane *et al*., 1987). We chose 100 µM as the working concentration of PCA as this is likely to be in a physiologically relevant range based on concentrations measured both in pure cultures and in the field. In broth cultures of biocontrol strains of *Pseudomonas*, PCA accumulates to concentrations ranging from dozens to hundreds of micromolar (Séveno *et al*., 2001; Tagele *et al*., 2019). In natural wheat rhizospheres, PCA has been detected at nanomolar concentrations (Mavrodi *et al*., 2012), but these bulk measurements almost certainly underestimate local concentrations at the micron scale given that bacteria colonize the rhizosphere in a patchy manner (Thomashow *et al*., 1990). Notably, PCA can accumulate in biofilms to concentrations 360-fold greater on a per volume basis compared to broth cultures (Séveno *et al*., 2001), suggesting that local concentrations of PCA in the rhizosphere, where biocontrol strains form robust biofilms (LeTourneau *et al*., 2018), may be orders of magnitude higher than the reported bulk values.

The screen was performed using 24-well plates and 0.1x tryptic soy agar (TSA) that was either left unadjusted (pH 7.3-7.5) or adjusted to pH 5.1. Each isolate was spotted onto agar in separate wells to prevent crosstalk and antagonism between the strains, and image analysis was used to derive quantitative information about the growth of each strain over the course of 7 days (with one timepoint per day). Because several strains among our 166 isolates appeared to be duplicates of each other based on 16S sequence and colony morphology, we restricted the screen to 108 strains that we judged likely to be unique (Table S1A); where multiple strains appeared identical, we chose a representative strain. We also included 30 strains obtained from public culture collections that represented species found among our isolates (Table S1B), in order to investigate the extent to which PCA resistance phenotypes are consistent within species across different strains isolated from different geographical locations. Accurate quantification of growth was not possible in this screen for certain strains that formed transparent colonies, produced dark pigments, or had a tendency to turn mucoid and spread (Fig. S2). Nevertheless, this assay enabled us to derive detailed profiles of PCA sensitivity and resistance for the vast majority of our strains.

To compare PCA susceptibility across strains, we first focused on a single-timepoint snapshot of each strain’s phenotype taken at the equivalent of early stationary phase (i.e. around the time that the spots on non-PCA control plates reached their maximal density). In accordance with previous reports of the pH dependency of PCA toxicity, we found that more of our strains were sensitive to PCA at pH 5.1 than at pH 7.3, and strains that were mildly inhibited by PCA at pH 7.3 were typically inhibited more strongly at pH 5.1 (Fig. 2A-B). At pH 5.1, strains of Actinobacteria and Firmicutes were strongly inhibited by PCA, while most Proteobacteria were relatively resistant. Members of Bacteroidetes exhibited variable responses, ranging from mild to severe growth inhibition (Fig. 2A-B). At pH 7.3, there was considerably more phenotypic variation among the Actinobacteria (Fig. 2A-B), particularly within the *Streptomyces* genus. In some cases, there was even noticeable variation within the same putative species of *Streptomyces* (Fig. S3). In addition, two strains of *Microbacterium* (W2I7 and W4I20), as well as one strain of *Arthrobacter* (W3I6), grew slightly better in the presence of PCA at pH 7.3 compared to the control condition, perhaps suggesting that they can use PCA as a carbon source or otherwise benefit from its presence at a pH where toxicity is limited (Fig. 2B). On the other hand, in contrast to the phenotypic variation seen among Actinobacteria, Firmicutes remained almost universally sensitive to PCA at pH 7.3 (albeit somewhat less so than at pH 5.1), while all members of Bacteroidetes were resistant to PCA at pH 7.3, in contrast to their sensitivity to PCA at pH 5.1 (Fig. 2A-B). Proteobacteria remained generally resistant (Fig. 2A-B), with the most notable exception being a strain of *Sphingomonas faeni*, isolate W4I17, that was inhibited by PCA at both pH 7.3 and pH 5.1 (Fig. 2A-B).

**Figure 2:**
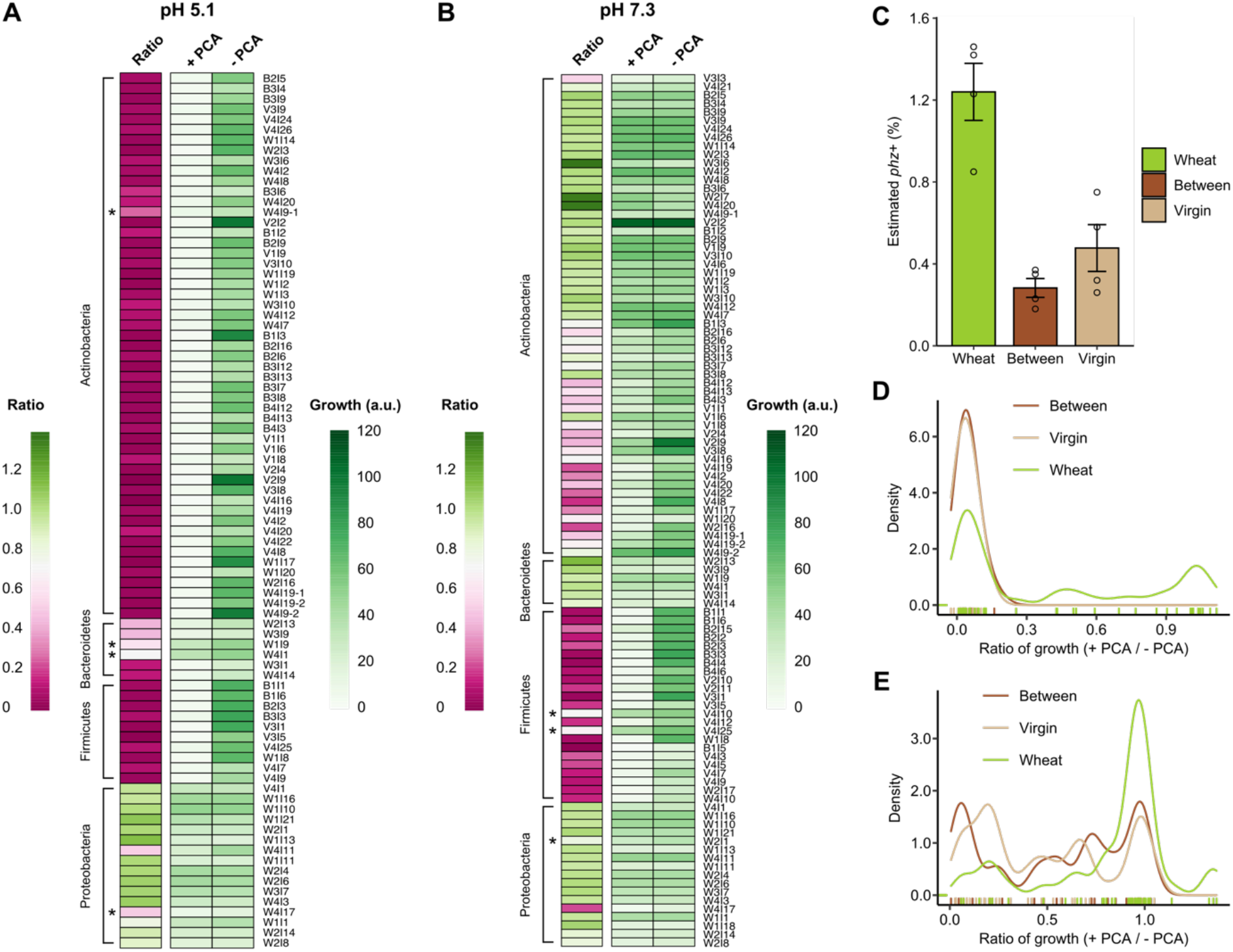
Distribution of PCA resistance phenotypes across phyla and soil sample type. **A-B**. Heat maps depicting the growth of the strains at pH 5.1 (A) and pH 7.3 (B). Each row represents a strain. The leftmost columns are colored according to the ratio of growth on PCA-containing agar versus PCA-free agar; magenta indicates sensitivity to PCA while green indicates resistance to PCA. The right two columns in each heat map are colored according to the separate values for growth on PCA-containing agar (+ PCA) or solvent control agar (-PCA), with darker green indicating more growth. Values are the mean of two to four biological replicates that each comprised three technical replicates. Asterisks indicate strains that displayed markedly variable susceptibility to PCA across biological replicates. Strains for which fewer than two biological replicates grew to stationary phase in the PCA-free control at the given pH, or for which color interfered with growth quantification, were omitted from this analysis. See Methods for a description of how growth was quantified. **C**. Relative abundance of putative phenazine producers (*phz*+ bacteria) across the three types of soil samples. Error bars represent the standard deviation. **D-E**. Density plots (i.e. smoothed histograms) representing the distribution of PCA resistance phenotypes at pH 5.1 (D) and pH 7.3 (E) among strains isolated from Between (bulk), Virgin (bulk), or Wheat (rhizosphere) soil. Higher values along the x-axis indicate greater resistance to PCA. If multiple identical strains were isolated from the same soil type, only one representative was counted; thus, 26 strains were isolated from Between, 39 from Virgin, and 45 from Wheat. Colored tick marks above the x-axis represent where individual isolates fall along the range of PCA resistance phenotypes.

Interestingly, while most strains yielded qualitatively consistent results across independent biological replicates that were separated by months (Fig. S4-S7), a handful of strains displayed variable growth and lag times at pH 5.1 either in the absence of PCA (mostly among Actinobacteria and Firmicutes), or in the presence of PCA (a few strains among the Bacteroidetes). Growth at pH 7.3 was generally more consistent, with the exception being two strains of *Neobacillus niacini* (V4I10 and V4I25) that initially failed to grow in the presence of PCA, but grew with minimal to no inhibition in subsequent replicates. The reasons for these discrepancies are unclear, but may be related to variations in how long the strains had been in stationary phase prior to inoculating the experimental cultures, which was difficult to control precisely due to differing growth rates across the large number of strains. The trace nutrient content may also have varied between different batches of media, as the first replicate was performed using a different lot of tryptic soy broth compared to subsequent replicates. Thus, the growth and PCA sensitivity of some strains may be influenced by environmental factors beyond pH that remain to be elucidated.

We next examined whether there was any evidence of correlation between PCA resistance phenotypes and the type of soil each strain was isolated from (Between, Virgin, or Wheat). A previous study based on samples taken from the same wheat field found that the relative abundance of phenazine producers was higher in the wheat rhizosphere compared to adjacent bulk soil (Dar *et al*., 2020), a finding that we also confirmed using shotgun metagenomic sequencing of our soil samples (Fig. 2C). We therefore hypothesized that if PCA-mediated antibiosis had shaped the bacterial community composition of this field, the prevalence of PCA resistance would be higher among isolates from the rhizosphere samples. Indeed, the Wheat samples clearly had the highest proportion of PCA-resistant isolates, regardless of whether resistance was assessed at pH 5.1 (Fig. 2D) or pH 7.3 (Fig. 2E). By contrast, all strains from Between or Virgin samples were highly sensitive to PCA at pH 5.1, though at pH 7.3, the resistance phenotypes of isolates from either type of bulk soil spanned the full range from highly sensitive to completely resistant. However, these findings are conflated with the fact that all of our Bacteroidetes strains, and all but one strain of Proteobacteria, were exclusively isolated from the Wheat samples, and that the members of these two phyla were generally relatively resistant to PCA, especially at pH 7.3. Further experiments will be necessary in order to distinguish whether the enrichment of PCA-resistant phenotypes in the rhizosphere samples is a consequence of PCA-mediated antibiosis or merely an indirect reflection of other factors that favor the growth of Proteobacteria and Bacteroidetes in the rhizosphere.

Finally, visualizing the growth of each strain over the full 7-day time course revealed more nuanced variations among the PCA resistance phenotypes at both pH 5.1 and pH 7.3 (Figs. 3 and S4-S7). For example, while some strains of Bacteroidetes displayed increased lag and/or slower growth rates in the presence of PCA at pH 5.1, the growth on PCA often eventually caught up to the PCA-free controls. At pH 7.3, the effect of PCA on *Peribacillus* species was a combination of increased lag and lower final cell density, but apparently not decreased maximal growth rate, as compared to no growth at all on PCA at pH 5.1. This was distinct from the slower growth rates seen for certain strains of *Bacillus, Neobacillus*, and *Paenibacillus* at pH 7.3. A few strains of the latter three genera appeared to be still unable to grow on PCA at pH 7.3 based on the values derived from image analysis (Fig. S5). However, examination of the plate images by eye revealed that this sometimes reflected a limitation in the sensitivity of our imaging and quantification methods to low levels of growth, rather than a true lack of growth (Fig. S2).

**Figure 3:**
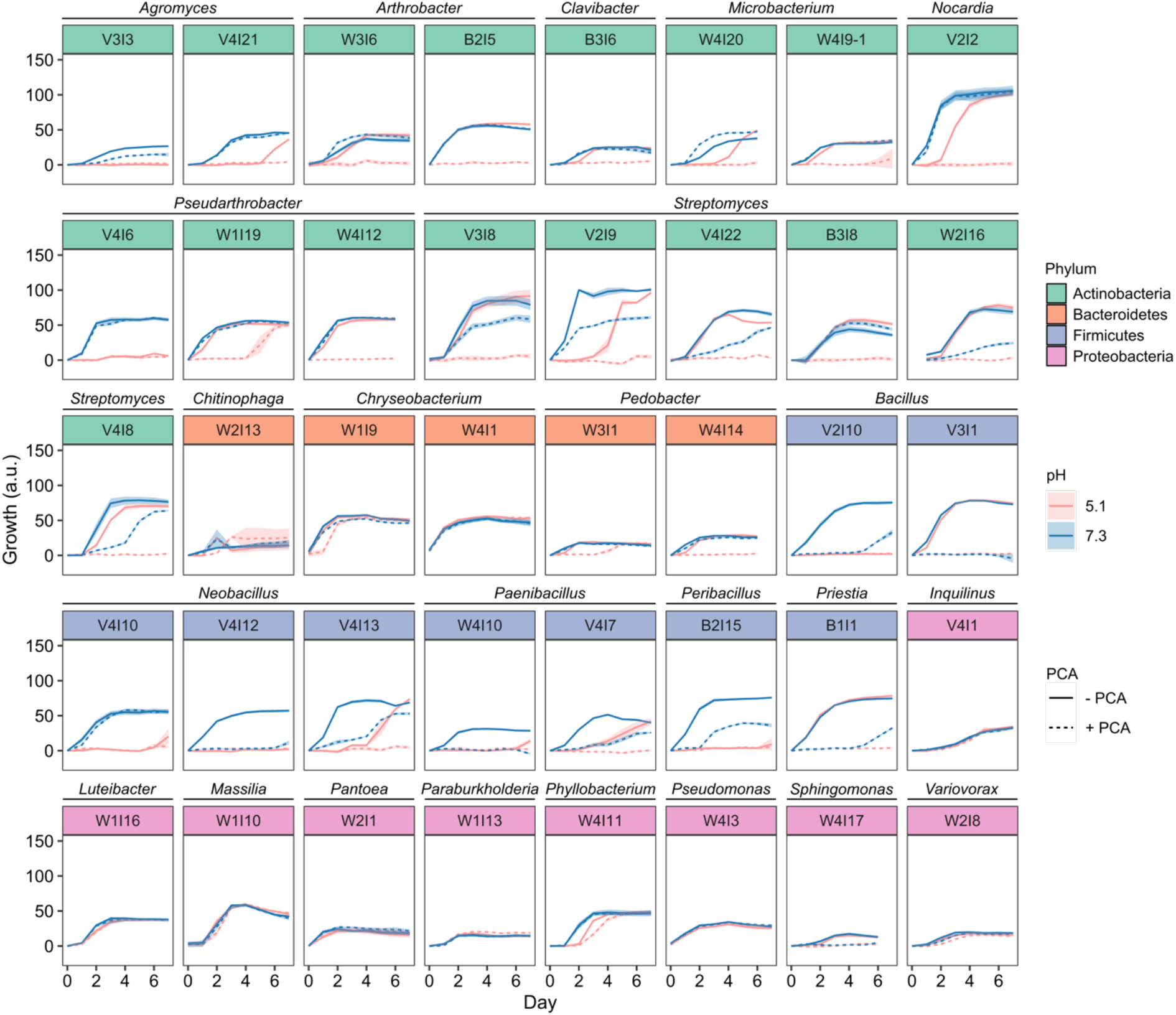
Growth of representative strains over time with and without PCA. Growth was quantified as described in the Methods. Solid lines represent the growth of spotted cultures on PCA-free agar, and dashed lines represent the growth of spotted cultures on agar containing 100 µM PCA. Blue represents growth at pH 7.3 and pink represents growth at pH 5.1. Data points are the mean of three technical replicates from a representative biological replicate for each strain, and the shaded ribbon represents the standard deviation. The selected strains depicted here cover the range of PCA susceptibility phenotypes observed in each genus. The full set of biological replicates for all tested isolates is displayed in supplementary Figs. S4-S7.

### PCA reduction rates in relation to PCA susceptibility

We next sought to determine whether there are physiological correlates of PCA susceptibility or resistance across taxonomically-diverse bacteria. Given that phenazines are redox-active molecules whose toxicity is thought to be related to ROS generation as a consequence of redox cycling in the cell (Hassan and Fridovich, 1980; Mavrodi *et al*., 2006), we hypothesized that susceptibility to PCA would be correlated with higher redox-cycling rates. PCA oxidation occurs rapidly and abiotically in the presence of oxygen (Wang and Newman, 2008), suggesting that the rate of PCA reduction would be the primary driver of differences in redox-cycling rates across strains under oxic conditions. Therefore, to test our hypothesis, we measured the rate of PCA reduction under anoxia as a proxy for the true redox-cycling rate, using a subset of strains that covered a range of taxonomic groups and PCA susceptibility phenotypes.

For each tested strain, the PCA reduction rate was typically higher at pH 5.1 than at pH 7.3 (Fig. 4A), matching the expectation that PCA uptake would tend to be faster at a lower pH and the observation that most susceptible strains were more sensitive to PCA at pH 5.1. The exceptions were: 1) strains belonging to the Firmicutes, which had lower reduction rates at pH 5.1 in our assay despite being more susceptible to PCA under this condition, and 2) *Sphingomonas faeni* W4I17, which was in fact more susceptible to PCA at pH 7.3 than at pH 5.1 (Fig. 2A-B), matching the pH dependency of its rate of PCA reduction. Notably, some of the tested Firmicutes often struggled to grow at pH 5.1 even in the absence of PCA (Fig. S5). This observation might account for the inhibitory effect of acidic pH on PCA reduction in these strains, given that a functional metabolism would be required for the generation of cellular reductants, such as NADH, that can indirectly or directly reduce PCA. For the strains of Firmicutes that were able to grow at acidic pH (namely B1I1 and B1I6), the reason for their lower PCA reduction rate at pH 5.1 remains unclear.

**Figure 4:**
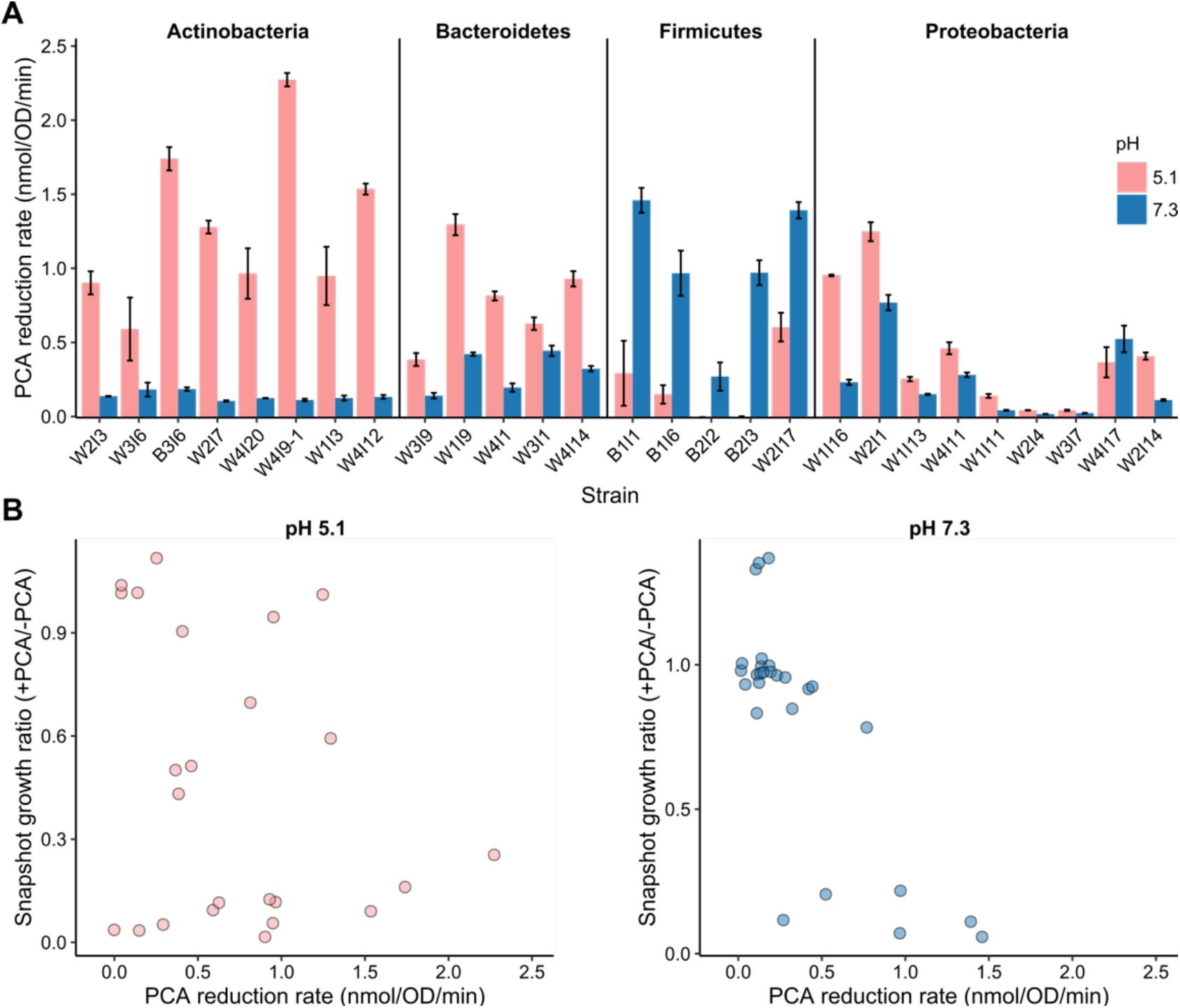
PCA reduction rates of selected strains at pH 5.1 and pH 7.3. **A**. PCA reduction rates of representative strains from each phylum at pH 5.1 and pH 7.3. The values represent the mean of three biological replicates and error bars represent the standard deviation. Reduction rates were normalized to OD_600_ values. **B**. PCA reduction rates versus each strain’s growth ratio (+PCA/-PCA) at early stationary phase. The growth ratios are the same values calculated and used in Fig. 2.

We also examined whether there was any correlation between PCA reduction rates and resistance levels across different strains at each pH. To do so, we plotted each strain’s PCA reduction rate against the early stationary phase snapshot ratio of growth on PCA versus no PCA (Fig. 4B). We also calculated Spearman’s correlation coefficient (*r*_*s*_). Interestingly, there was a statistically significant negative correlation between reduction rate and resistance at pH 7.3 (*r*_*s*_ = −0.70, *p* < 0.001), but not at pH 5.1 (*r*_*s*_ = −0.16, *p* = 0.4485). However, there were exceptions to the trend even at pH 7.3; for example, while PCA severely and consistently inhibited the growth of the four fastest PCA-reducing strains, the fifth-fastest strain, W2I1 (*Pantoea agglomerans*), appeared to be completely resistant to PCA in one biological replicate, though its growth was mildly inhibited by PCA in another replicate (Fig. S7). In addition, strains with reduction rates in the middle range of 0.2-0.4 nmol/OD_600_/min spanned the full range of resistance phenotypes, from a growth ratio of 0.12 (*Peribacillus simplex* B2I2) to 0.96 (*Phyllobacterium ifrigiyense* W4I11). Taken together, these data suggest that a low reduction rate may be helpful in mitigating PCA toxicity but is neither necessary nor sufficient for resistance.

### Effects of efflux pump inhibitors on PCA susceptibility

Given that PCA reduction rate alone was insufficient to account for the range of sensitivities to PCA, we asked if efflux pumps might serve as an important contributor to PCA resistance in some strains. To address this question, we used two efflux pump inhibitors (EPIs) in combination: 1) reserpine, which is thought to target efflux pumps in the major facilitator superfamily (MFS) (Kumar *et al*., 2013), and 2) phenylalanine-arginine β-naphthylamide (PAβN), which is thought to target resistance-nodulation-division (RND) efflux pumps (Jamshidi *et al*., 2017), although it may also permeabilize the outer membrane of Gram-negative bacteria (Lamers *et al*., 2013; Schuster *et al*., 2019). We tested these EPIs on a subset of the strains that were used in the PCA reduction assay; we excluded the *Bacillus* strains due to their tendency to settle on the bottom of the wells during growth in the plate reader, which confounded the OD_600_ readings. Preliminary experiments revealed that different strains were differentially sensitive to the EPIs themselves, with growth inhibition evident even in PCA-free controls. We therefore determined for each strain the maximal EPI concentration that, when the two EPIs were used separately, did not markedly reduce growth of PCA-free controls (up to a maximum dose of 20 µg/mL for reserpine and 50 µg/mL for PAβN) (Table S2). Interestingly, for many strains, the combination of both EPIs remained highly toxic even after this optimization step, particularly at pH 7.3, possibly indicating synergistic toxicity or redundant roles of different efflux pumps in preventing intracellular accumulation of toxic metabolic intermediates (Fig. S9); the toxicity appeared to be pH-dependent, suggesting that the protonation state of the EPIs might influence their efficacy. In these cases, it was not possible to assess the impact of efflux pump inhibition on PCA sensitivity. However, for the remaining strains, three types of responses to the EPIs emerged. For one group of strains, treatment with both EPIs in combination did not affect their resistance to PCA (Fig. 5A-B); however, we cannot distinguish whether this is because efflux does not contribute to PCA resistance in these strains, or because the tested EPIs did not effectively target the relevant efflux pumps. For the second group of strains, the EPIs enhanced the toxicity of PCA, as would be expected if efflux is an important mechanism of resistance (Fig. 5C). Finally, for the third group of strains, the EPIs counterintuitively appeared to mitigate the toxicity of PCA, or vice versa (Fig. 5D). Although unexpected, the latter phenomenon indirectly suggests that PCA interacts with efflux systems in these strains. For example, treatment with either PCA or the EPIs might upregulate the expression of certain efflux pumps, leading to decreased toxicity relative to treatment with either type of toxin alone. Alternatively, given that some pumps in the MFS family are thought to be bidirectional (Pao *et al*., 1998), it is possible that MFS pumps promote uptake rather than export of PCA in certain strains.

**Figure 5:**
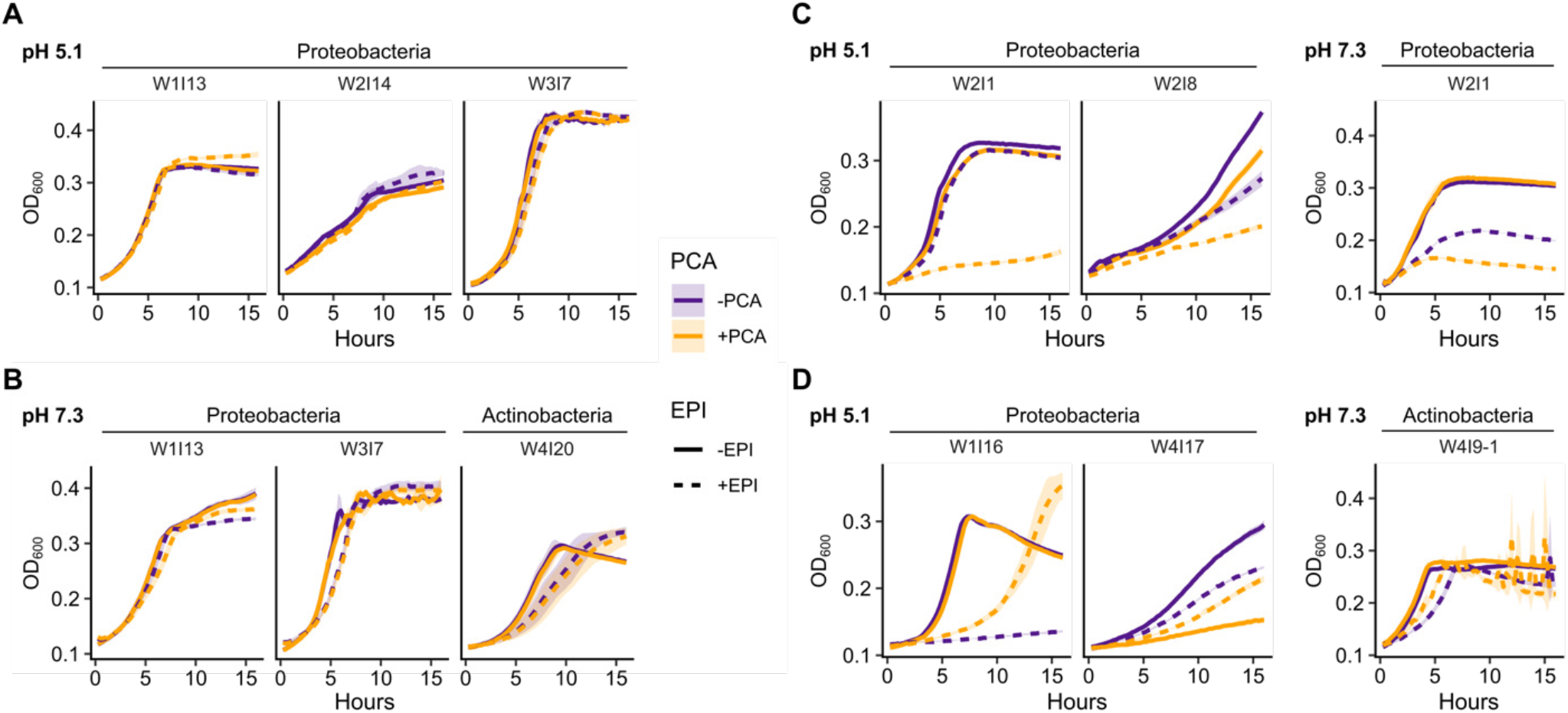
Effects of efflux pump inhibitors on PCA susceptibility. **A-B**. Growth curves of strains for which treatment with a combination of the efflux pump inhibitors (EPI) reserpine and PAβN did not affect susceptibility to PCA at pH 5.1 (A) or pH 7.3 (B). **C**. Growth curves of strains for which treatment with reserpine and PAβN potentiated the toxicity of PCA. **D**. Growth curves of strains for which treatment with reserpine and PAβN improved growth in the presence of PCA (strain W4I17) or vice versa (strains W1I16 and W4I9-1). In all panels, data points are the mean of three biological replicates and the shaded ribbon represents the standard deviation. The concentrations of reserpine and PAβN used for each strain can be found in Table S2.

## Discussion

In this study, we have characterized profiles of resistance to an agriculturally-relevant phenazine across taxonomically diverse bacteria isolated from a wheat field where phenazine producers are indigenous. We also examined potential physiological correlates of phenazine resistance in a subset of these strains. Our findings establish a basis for inferring whether intrinsic resistance is a factor that affects how phenazine production shapes bacterial communities in the rhizosphere. In particular, it will now be possible to test data-driven predictions regarding which strains, species, or even phyla are most likely to be affected by phenazine-mediated antibiosis, even though the mechanistic underpinnings of phenazine susceptibility or resistance are likely multifactorial and appear to differ across species.

One of our more notable findings is that the previously reported pH dependency of PCA toxicity varies across bacterial taxonomic groups. For most Actinobacteria and Firmicutes, sensitivity to PCA was clearly higher at pH 5.1 than at pH 7.3, indicating greater toxicity at the lower pH as expected according to the pKa of PCA. On the other hand, nearly all tested Proteobacteria were completely resistant to 100 µM PCA regardless of pH, at least down to pH 5.1. Consequently, at pH 5.1, PCA resistance phenotypes largely correlated with phylum—and more broadly, a Gram-positive versus Gram-negative divide, which has previously been reported for the toxicity of another phenazine, pyocyanin (Baron and Rowe, 1981). The Gram-positive versus Gram-negative divide at pH 5.1 is perhaps not surprising, as at this pH, a significant proportion (∼14%) of PCA in solution is protonated and therefore would not be repelled by the negatively charged cell wall. Under this condition, the outer membrane of Gram-negative species presumably presents an additional barrier to the entry of PCA, helping to limit the intracellular accumulation of the toxin in the same manner as for numerous other antibiotics (Nikaido, 1989). Less expected, however, was the within-phylum and even within-species variation in PCA resistance phenotypes at pH 7.3 among certain taxonomic groups. Interestingly, several *Streptomyces* strains exhibited at least some PCA-dependent growth inhibition at pH 7.3, even though many *Streptomyces* species are capable of producing their own toxic phenazines (Turner and Messenger, 1986; Dar *et al*., 2020). Given that the major limit on PCA toxicity at circumneutral pH is thought to be its ability to enter cells, the phenotypic variability at pH 7.3 may indicate the presence of transporters or channels capable of phenazine uptake in some PCA-sensitive Actinobacteria. Alternatively, it is possible that some PCA-sensitive strains locally acidified the growth medium, which was only weakly buffered (1.4 mM K_2_HPO_4_).

Importantly, despite the general Gram-positive versus Gram-negative divide in sensitivity to PCA at pH 5.1, possessing an outer membrane is evidently not a leakproof shield against PCA toxicity. Proteobacteria as a group were more resistant to PCA at pH 5.1 compared to strains of Bacteroidetes, most of which exhibited increased lag in the presence of PCA, even though both phyla comprise Gram-negative bacteria. Intriguingly, one major difference between these clades is that Proteobacteria generally utilize ubiquinone as an electron carrier during aerobic growth (Collins and Jones, 1981), while the Bacteroidetes genera screened in this study (*Chitinophaga, Chryseobacterium*, and *Pedobacter*) utilize menaquinone (Lin *et al*., 2015; Singh *et al*., 2017; Kong *et al*., 2019). Menaquinones have a lower reduction potential compared to ubiquinones (White *et al*., 2012). Although the standard reduction potential of menaquinone is still higher than that of PCA (−74 mV compared to −177 mV) (Price-Whelan *et al*., 2006; White *et al*., 2012), indicating that PCA likely is not reduced by menaquinol, this difference with ubiquinone nevertheless raises the possibility that PCA may interact differently, and perhaps more readily, with the aerobic respiratory electron transport chain of menaquinone-utilizing Bacteroidetes strains compared to Proteobacteria, thereby generating more ROS and/or interfering with the generation of ATP. Interestingly, another study has shown that the reduced form of different phenazine with a low reduction potential, neutral red (3-amino-7-dimethylamino-2-methylphenazine), can directly transfer electrons to menaquinone, bypassing the proton-pumping NADH dehydrogenase complex that normally transfers electrons from NADH to menaquinone and thereby “short-circuiting” the electron transport chain. We hypothesize that a similar phenomenon may occur with PCA in strains that rely on menaquinone. In future studies, this hypothesis could be tested by 1) performing *in vitro* experiments with purified menaquinone, ubiquinone, and reduced PCA to directly test whether quinones oxidize the latter and if so, whether the kinetics differ between menaquinone and ubiquinone, 2) measuring ROS production and steady-state ATP pools in selected strains of Bacteroidetes and Proteobacteria both in the presence and absence of PCA, and 3) forcing species of Proteobacteria (such as *Escherichia coli*) that utilize both ubiquinone and menaquinone in different branches of their electron transport chains to rely only on the latter (for example, by deleting the genes for ubiquinone biosynthesis), followed by reassessing their sensitivity to PCA to see if there is any effect.

Beyond the specific strains screened in this study, the risk assessment of phenazine-producing biocontrol strains, and the understanding of phenazine biology in general, would benefit greatly from the development of a platform for the prediction of phenazine resistance phenotypes from genomic or phylogenetic information. The results of this study already indicate that, depending on the environmental pH, phylogenetic information may be of limited utility for prediction of PCA resistance, given the variation in phenotypes for Actinobacteria at circumneutral pH. However, this variability may also hold the key to identifying genome-based predictive markers of phenazine resistance. Given the taxonomic and phenotypic diversity of our strain collection, whole-genome sequencing of our isolates, followed by comparative genomics, could potentially reveal such markers.

In summary, this work has laid the groundwork for rectifying a major gap in studies of how introduction of a phenazine-producing biocontrol strain affects rhizosphere bacterial communities. Previous studies have lacked information about the baseline prevalence of resistance in the native communities (Gagliardi *et al*., 2001; Bankhead *et al*., 2004; Kozdrój *et al*., 2004; Roquigny *et al*., 2018), and in the absence of such information, it is impossible to determine whether a negative result (lack of change in the rhizosphere community) reflects a high prevalence of resistance to PCA that is particular to the studied community, versus a general lack of toxicity of PCA to most bacteria or fundamental abiotic constraints on the antibacterial activity of PCA in the rhizosphere (e.g. limited diffusion, adsorption to soil particles, etc.). Distinguishing between these scenarios is key to assessing the risk of unwanted side effects in rhizosphere communities upon the application of phenazine-producing biocontrol strains. In addition, recent studies have demonstrated that phenazines produced by the opportunistic pathogen *Pseudomonas aeruginosa* can promote bacterial tolerance and resistance to clinical antibiotics (Schiessl *et al*., 2019; Zhu *et al*., 2019; Meirelles *et al*., 2021; VanDrisse *et al*., 2021), and that these effects can extend to other opportunistic pathogens that are resistant to phenazines (Meirelles *et al*., 2021). Thus, understanding the prevalence and genetic determinants of resistance to phenazines may have implications not only for agriculture but also for human medicine and beyond, as we continue to uncover new ecological roles for these multifaceted bacterial metabolites.

## Methods

### Isolation of bacteria from wheat rhizosphere and bulk soil samples

Three types of samples were collected from a non-irrigated wheat field at Washington State University’s Lind Dryland Research Station on August 9, 2019: wheat plants and surrounding soil, bulk soil from in between the planted rows, and bulk soil from a “virgin” hillside site that has never been farmed. The wheat had been harvested a few weeks prior to sample collection. All samples were immediately stored on ice in clean plastic bags, and subsequently at 4 °C for four days until processing. Rhizosphere soil samples were obtained by shaking the wheat roots until only 1-2 mm of tightly-adhering soil remained, followed by excising the roots at the crown with a sterile razor blade. The roots of 2-3 plants per replicate were placed in 50 mL conical tubes with 30 mL of sterile deionized water, vortexed at top speed for 1 min, and treated in an ultrasonic water bath for 1 min to dislodge bacteria from the roots. Bulk soil samples (1 g per sample) were processed in the same manner. Large soil particles were allowed to settle to the bottom of the tubes on the bench top, and 100 µL each of a 10-fold dilution series of the supernatants was spread onto 0.1x TSA plates containing 50 µg/mL nystatin to inhibit fungal growth. The plates were incubated upside down at room temperature in the dark and monitored for the appearance of new colonies over the course of a week. Colonies that appeared morphologically distinct in each sample were picked and restreaked on 0.1x TSA until visually pure cultures were obtained. Multiple representatives were also picked for the most common colony types in an attempt to account for strain variations that might not be apparent to the eye. Once the streaks yielded uniform single colonies, the isolates were inoculated into 1.5 mL of 0.1x tryptic soy broth (TSB) in 5 mL polycarbonate culture tubes and incubated at 30 °C with shaking at 250 rpm. After 1-3 days of incubation, depending on when the cultures became turbid, 0.5 mL of each culture was mixed with 0.5 mL of 50% glycerol and stored at −80 °C. Some cultures never became turbid under these growth conditions, but nevertheless yielded viable frozen stocks.

### Species identification by 16S rRNA gene sequencing

Single colonies or patches of morphologically pure streaks were picked and resuspended in 20 µL sterile nuclease-free water. Colony PCR was performed using GoTaq Green Master Mix (Promega) in 50 µL reactions (1 µL of cell suspension) according to the manufacturer’s instructions. For putative streptomycetes (isolates that formed hard colonies rooted in agar, often with aerial hyphae), the thermocycling protocol was modified to include a 10 min initial heating step at 95 °C (compared to 2 minutes for other samples). The primers used were 27F (AGAGTTTGATCMTGGCTCAG) and 1492R (TACGGYTACCTTGTTACGACTT) (Lane, 1991). The PCR products were run on a 1% agarose gel to verify the presence of a single band at the expected size (∼1500 bp), followed by purification with the Monarch PCR and DNA Cleanup Kit (New England Biolabs). The purified products were submitted for Sanger sequencing at Laragen, Inc., using the same 27F and 1492R primers. The resulting forward and reverse sequences were aligned using MAFFT (https://mafft.cbrc.jp/alignment/server/) and subjected to BLAST against the NCBI 16S ribosomal RNA sequences database. For a few strains, full-length sequences could only be obtained following PCR on genomic DNA extracted using the DNeasy Blood & Tissue kit (Qiagen), following the manufacturer’s instructions for Gram-positive bacteria. In addition, for two strains (B1I5 and B2I8), the forward sequence consistently appeared to contain multiple products. We presumed that this was due to multiple primer binding sites or another sequencing artifact rather than mixed cultures as the corresponding sequences from the other direction were clean. In these cases, only the clean sequence was submitted to BLAST.

### PCA resistance screen

The optimized version of the PCA resistance screen was performed with four conditions: 0.1x TSA (15 g/L agar plus 3 g/L tryptic soy broth no. 2 from MilliporeSigma) with or without 100 µM PCA at pH 7.3 or pH 5.1 (adjusted with HCl). PCA was purchased from Princeton BioMolecular Research and dissolved in filter-sterilized 14 mM NaOH to make 10 mM stock solutions. The PCA stock solution or solvent control (14 mM NaOH) was added at 1% v/v to autoclaved molten 0.1x TSA; we verified that this addition did not noticeably alter the pH of the medium by pipetting aliquots onto pH test strips. Subsequently, 1 mL of the medium was pipetted into each well in 24-well polystyrene Cellstar Cell Culture plates (Greiner Bio-One). The plates were allowed to set and dry with the lids off in a biological safety cabinet for 20-30 min, followed by storage upside with the lids on at room temperature in the dark for two days prior to use.

Cell suspensions for inoculation in the screen were prepared in one of two ways. First, individual strains were inoculated into 5 mL TSB cultures in glass culture tubes and incubated at 25 °C with shaking at 250 rpm. Strains that grew overnight were then diluted to an OD_600_ of 0.05. Certain strains did not grow well in this condition—in particular, strains of *Agromyces, Streptomyces, Paenibacillus*, and *Neobacillus*. For these, some replicates were prepared by directly scraping cells from streaks grown on 0.1x TSA plates and resuspended the cells in 200 µL TSB with pipetting and brief vortexing at top speed. The OD_600_ of these cell suspensions was adjusted to 0.05, except for *Streptomyces* due to their filamentous nature and tendency to clump; instead, visible cell clumps were allowed to settle to the bottom and inocula were taken from the visually clear upper portion. Subsequently, 10 µL of each cell suspension was pipetted onto the agar in a single well in the 24-well plates. Three adjacent wells per condition were inoculated with each cell suspension, representing technical replicates. After the spots dried, the plates were incubated at room temperature upside in the dark for up to 7 days. Every 24 hrs, the plates were imaged in color at 600 dpi with an Epson Perfection V550 Photo flatbed scanner. For strains that were prepared both from liquid cultures and by resuspension from plates, we did not observe any consistent effect on the resulting PCA susceptibility phenotypes.

### Image analysis and quantification of growth

Images from the scanner were analyzed using Fiji (Schindelin *et al*., 2012). Circular regions of interest (ROIs) were drawn around each culture spot and the mean gray value was measured for each ROI. We also measured the mean gray values of equivalent ROIs in the wells of blank, uninoculated plates for each condition. The latter values were averaged across each 24-well blank plate to give the “background” gray value, which was then subtracted from the mean gray values of the culture spots. The resulting numbers were reported as the metric for growth. Importantly, while this method generally worked well for comparisons across conditions within each strain, there are a few caveats. First, this metric underestimated growth for strains that produced a dark pigment. Second, growth was difficult to quantify for a few strains that grew as nearly transparent colonies. Finally, this metric is not very sensitive to low levels of growth. Nevertheless, for the vast majority of strains, this approach captured visible differences in growth across the four conditions in the screen.

### Metagenomics-based estimation of the relative abundance of PCA producers

DNA was extracted from 250 mg of each soil sample using the DNeasy PowerSoil kit (Qiagen). Illumina metagenomic DNA libraries were prepared and sequenced (150 bp single-end reads) at the Millard and Muriel Jacobs Genetics and Genomics Laboratory at the California Institute of Technology. The resulting sequence data was processed according to the pipeline described by Dar et al. (2020).

### PCA reduction assay

Selected strains were grown in triplicate overnight in 5 mL TSB cultures at 25 °C with shaking at 250 rpm. The cells were then concentrated by serially centrifuging 1 mL aliquots at 9000 g for 2 min in microcentrifuge tubes, washed with 0.1x TSB (pH 7.3), and standardized to OD_600_ = 8 in 0.1x TSB (pH 7.3). Subsequently, 100 µL of each cell suspension was transferred into a 96-well plate, and the plate was moved into a Coy anaerobic chamber (5% hydrogen/95% nitrogen headspace). Using plasticware and media that had been passively degassed in the chamber for at least 3 days, electrochemically-reduced PCA was added to a final concentration of 100 µM in 190 µL of 0.1x TSB (pH 5.1 or pH 7.3) per well in a black clear-bottom 96-well plate; a standard curve was also prepared using reduced PCA at 1 µM, 10 µM, 50 µM, and 100 µM. Cells were added to each well (except the standard curve) to an OD_600_ of ∼0.4, and the plate was immediately transferred to a Biotek Synergy 5 plate reader maintained at 30 °C. OD_600_ and fluorescence readings (360 nm ex/528 nm em) were taken every 5 minutes, with continuous shaking on the “medium” setting. Reduction rates were derived from the slope of a linear regression fit to the visually linear range of the data using the function lm in R (see Fig. S8 for an example). The rates were converted to nmol/OD/min by relating the fluorescence units to the standard curve of reduced PCA concentrations and then dividing by the initial OD_600_ reading for each replicate. No appreciable change in OD_600_ was observed over the course of the experiment for any of the tested strains.

### Efflux pump inhibition growth curves

Reserpine was purchased from Sigma-Aldrich and dissolved in dimethyl sulfoxide (DMSO) to make a 20 mg/mL stock solution. PAβN was purchased from MedChemExpress and dissolved in distilled water to make a 50 mg/mL stock solution. Selected strains were grown in triplicate overnight in 5 mL TSB cultures at 25 °C with shaking at 250 rpm, washed once with 0.1x TSB (pH 7.3), and then inoculated to an initial OD_600_ of 0.05 in 160 µL of 0.1x TSB (pH 5.1 or pH 7.3) in a 96-well plate. Each well contained either a combination of reserpine and PAβN (see Table S2 for the concentrations used for each strain) or a solvent control (DMSO at 0.1% final concentration), along with either 0 or 100 µM PCA. The wells were topped with 50 µL of autoclaved mineral oil to prevent evaporation, and growth was monitored with OD_600_ readings every 15 minutes in a Biotek Synergy 5 plate reader set to 30 °C with continuous shaking on the “medium” setting.

## Data availability

The metagenomic sequencing data generated in this study were deposited in the Sequence Read Archive (SRA) under accession PRJNA521160.

## Acknowledgements

We thank Linda Thomashow and David Weller of the USDA Agricultural Research Service for providing the wheat rhizosphere and soil samples used in this study, and the members of the Newman lab for helpful feedback throughout the process of designing and analyzing the results of the PCA resistance screen. We also thank Zevin Condiotte for assistance with the initial rounds of testing the design of the PCA resistance screen and Richard Horak for assistance with extracting genomic DNA. This material is based upon work supported by the National Science Foundation Graduate Research Fellowship under Grant No. DGE-1745301. This work was also supported by grants to DKN from the ARO (W911NF-17-1-0024), NIH (1R01AI127850-01A1) and the Resnick Sustainability Institute at Caltech.

## Notes

### Competing Interest Statement

The authors have declared no competing interest.

### Summary of Updates

We have added the supplementary files, which were not automatically deposited at the time of initial submission.

