## Supplemental Figures 1-6 for "Prevalence and correlates of phenazine resistance in culturable bacteria from a dryland wheat field"

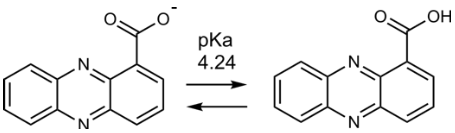


**Figure S1: Structures of protonated and deprotonated oxidized PCA.**
The negative charge on deprotonated PCA is thought to make PCA less toxic at higher pH by decreasing uptake due to the negative charge on cell membranes.

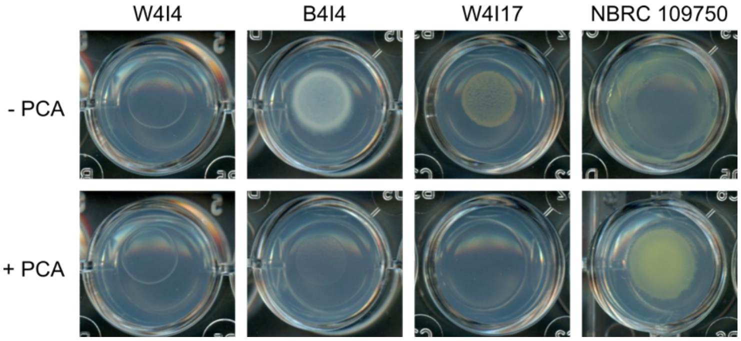


**Figure S2: Examples of image processing limitations for growth quantification.**
Accurate quantification of growth and PCA susceptibility was hindered in cases where the strains were transparent (e.g. *Microbacterium sp.* W4I4), grew to very low levels (e.g. *Neobacillus drentensis* B4I4 in the presence of PCA), produced pigments (e.g. *Sphingomonas faeni* W4I17), or had a tendency to turn mucoid and spread across the agar (*Chitinophaga ginsengisegetis* NBRC 109750). Such characteristics generally led to underestimation of growth. The displayed examples were grown at pH 7.3.

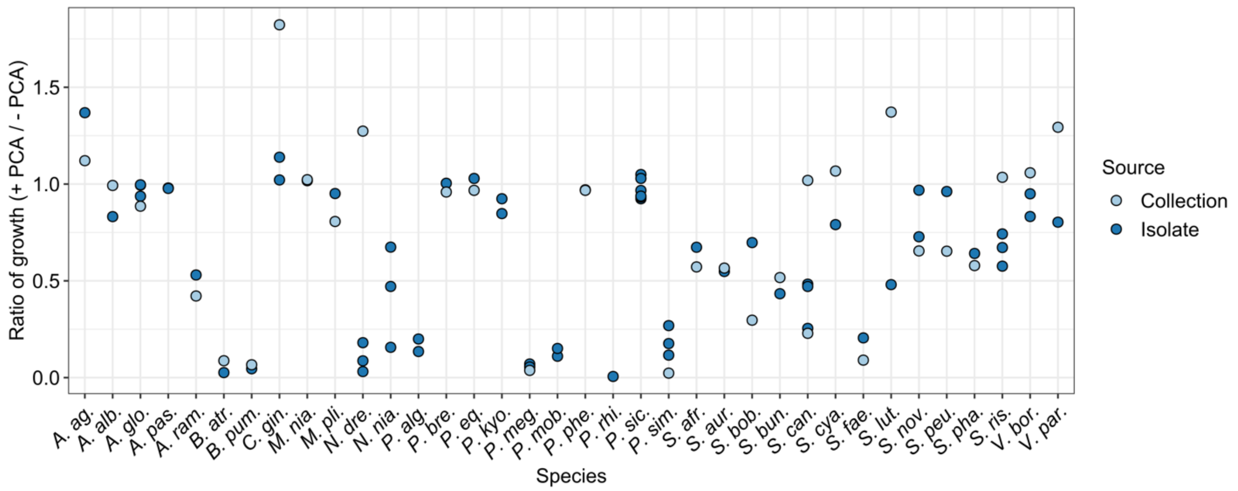


**Figure S3: PCA sensitivity at pH 7.3 of different strains belonging to the same putative species.** Dots represent the mean of two to four biological replicates for each strain (except for *S. phaeochromogenes* B-1248 and *S. bobili* B-1338, for which only one replicate was available). Dots are colored according to the source of the strain; dark blue represents wheat field isolates while light blue represents strains procured from public culture collections. The high ratio for the culture collection strain of *C. ginsengisegetis* reflects its tendency to become mucoid and spread while growing agar, which PCA appeared to restrict (see Fig. S2). *A. ag.* = *Arthrobacter agilis*; *A. alb.* = *Agromyces albus*; *A. glo.* = *Arthrobacter globiformus*; *A. pas.* = *Arthrobacter pascens*; *A. ram.* = *Agromyces ramosus*; *B. atr.* = *Bacillus atrophaeus*; *B. pum.* = *B. pumilus*; *C. gin.* = *Chitinophaga ginsengisegetis*; *M. nia.* = *Massilia niastensis*; *M. pli.* = *Massilia plicata*; *N. dre.* = *Neobacillus drentensis*; *N. nia.* = *Neobacillus niacini*; *P. alg.* = *Paenibacillus alginolyticus*; *P. bre.* = *Pseudomonas brenneri*; *P. eq.* = *Pseudarthrobacter equi*; *P. kyo.* = *Pedobacter kyongii*; *P. meg.* = *Priestia megaterium*; *P. mob.* = *Paenibacillus mobilis*; *P. phe.* = *Pseudarthrobacter phenanthrenivorans*; *P. rhi.* = *Paenibacillus rhizoryzae*; *P. sic.* = *Pseudarthrobacter siccitolerans*; *P. sim.* = *Peribacillus simplex*; *S. afr.* = *Streptomyces africanus*; *S. aur.* = *Streptomyces aurantiacus*; *S. bob.* = *Streptomyces bobili*; *S. can.* = *Streptomyces canus* (synonymous with *S. ciscaucasicus*); *S. cya.* = *Streptomyces cyaneofuscatus*; *S. fae.* = *Sphingomonas faeni*; *S. lut.* = *Streptomyces luteogriseus*; *S. nov.* = *S. novaecaesareae*; *S. peu.* = *Streptomyces peucetius*; *S. pha.* = *Streptomyces phaeochromogenes*; *S. ris.* = *Streptomyces rishiriensis*; *V. bor.* = *Variovorax boronicumulans*; *V. par.* = *Variovorax paradoxus*.

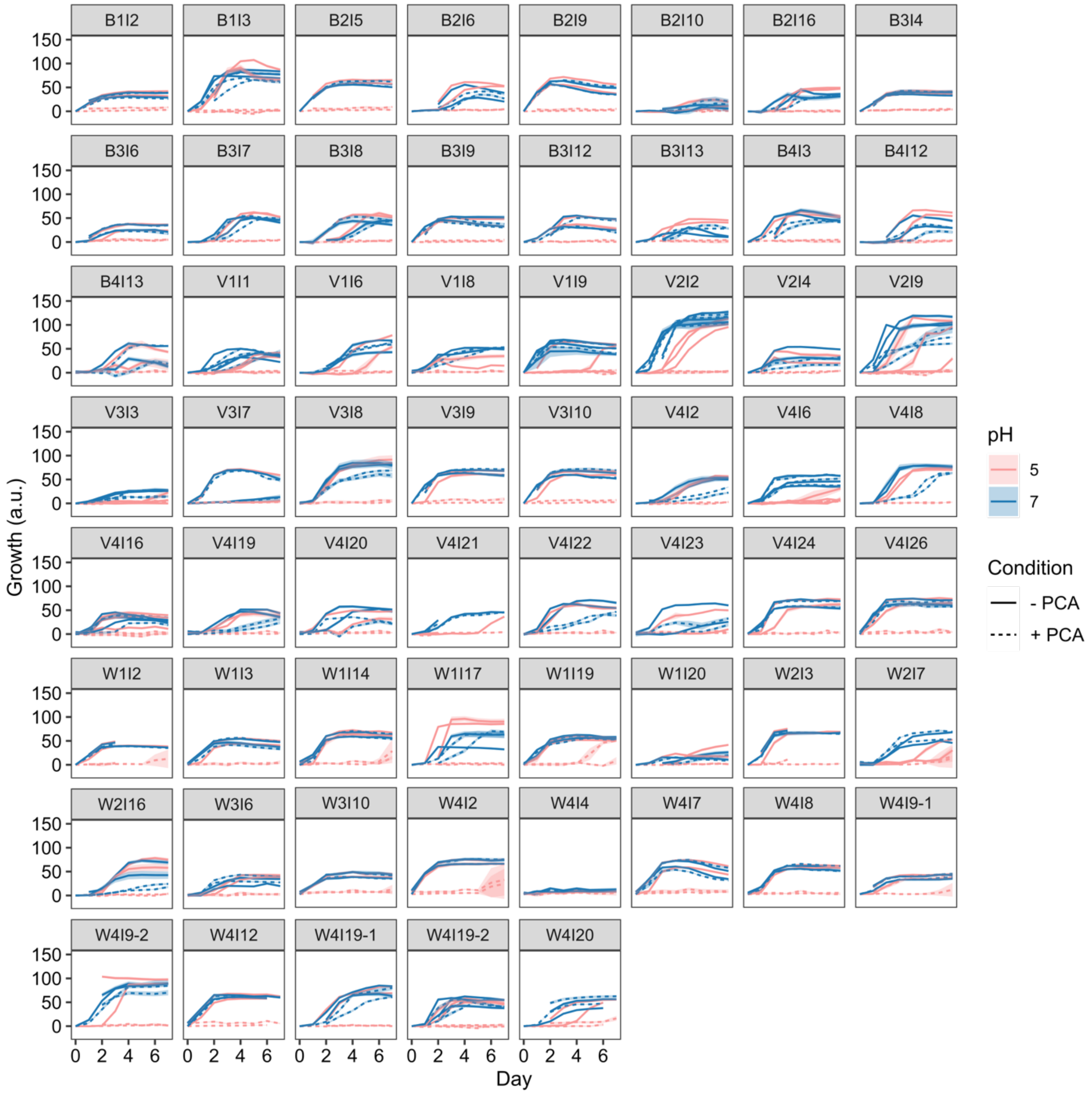


**Figure S4: Growth of Actinobacteria strains over time with and without PCA.**
Growth was quantified as described in the Methods. Solid lines represent the growth of spotted cultures on PCA-free agar, and dashed lines represent the growth of spotted cultures on agar containing 100 µM PCA. Blue represents growth at pH 7.3 and pink represents growth at pH 5.1. Data points are the mean of three technical replicates from a representative biological replicate for each strain, and the shaded ribbon represents the standard deviation. Different lines of the same color and line type (dashed vs. solid) represent separate biological replicates. For some biological replicates, data were collected only for a subset of days.

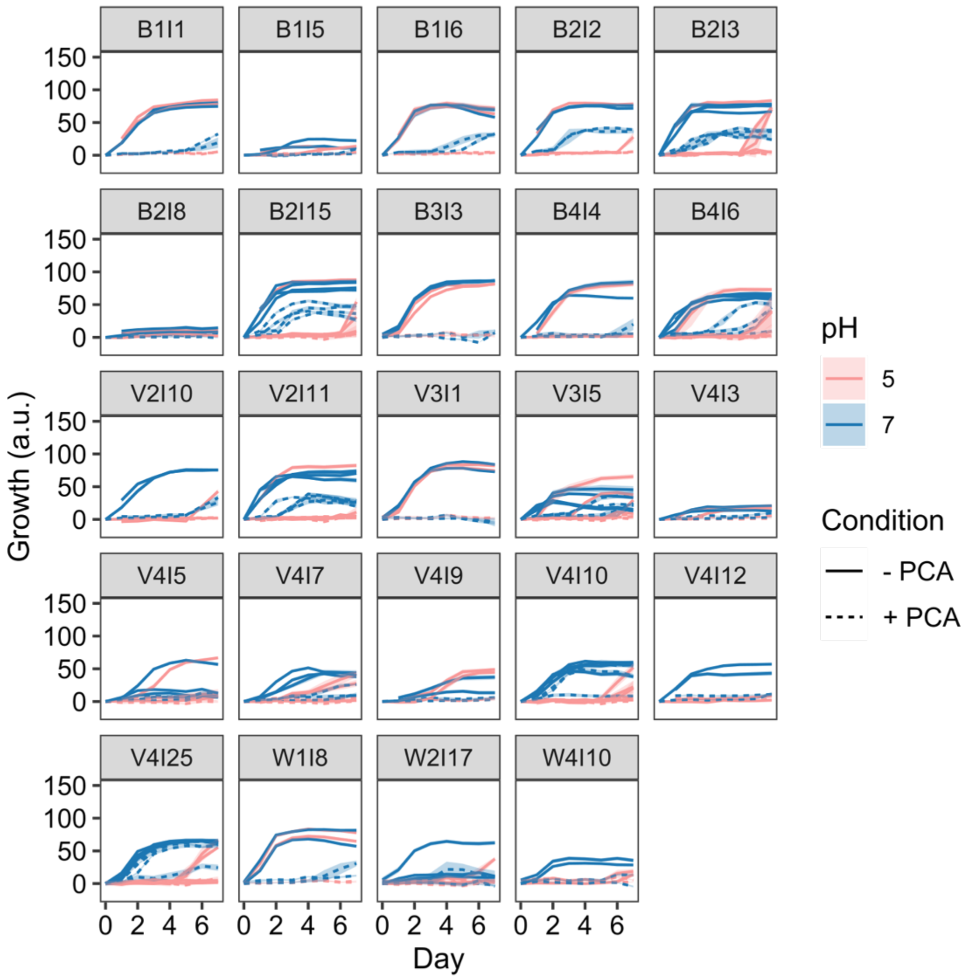


**Figure S5: Growth of Firmicutes strains over time with and without PCA.**
Growth was quantified as described in the Methods. Solid lines represent the growth of spotted cultures on PCA-free agar, and dashed lines represent the growth of spotted cultures on agar containing 100 µM PCA. Blue represents growth at pH 7.3 and pink represents growth at pH 5.1. Data points are the mean of three technical replicates from a representative biological replicate for each strain, and the shaded ribbon represents the standard deviation. Different lines of the same color and line type (dashed vs. solid) represent separate biological replicates. For some biological replicates, data were collected only for a subset of days.

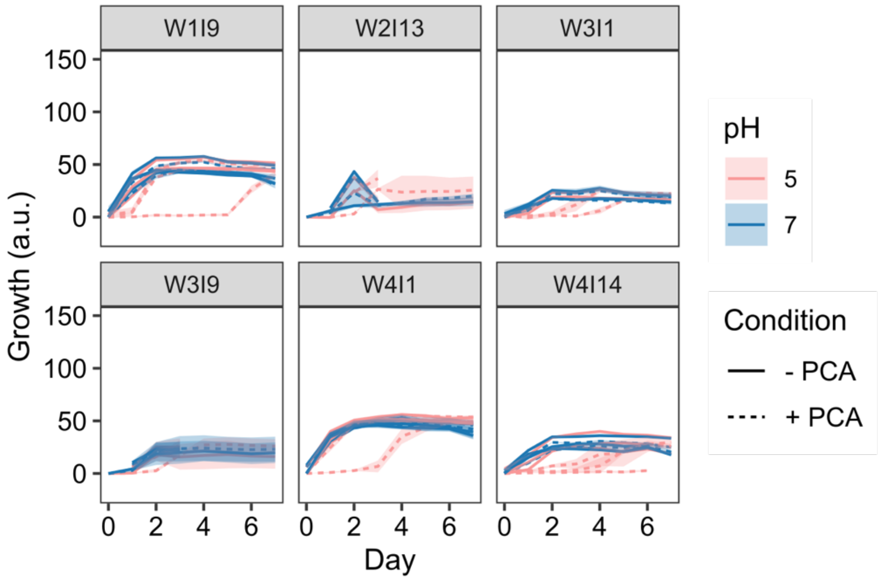


**Figure S6: Growth of Bacteroidetes strains over time with and without PCA.**
Growth was quantified as described in the Methods. Solid lines represent the growth of spotted cultures on PCA-free agar, and dashed lines represent the growth of spotted cultures on agar containing 100 µM PCA. Blue represents growth at pH 7.3 and pink represents growth at pH 5.1. Data points are the mean of three technical replicates from a representative biological replicate for each strain, and the shaded ribbon represents the standard deviation. Different lines of the same color and line type (dashed vs. solid) represent separate biological replicates. For some biological replicates, data were collected only for a subset of days.

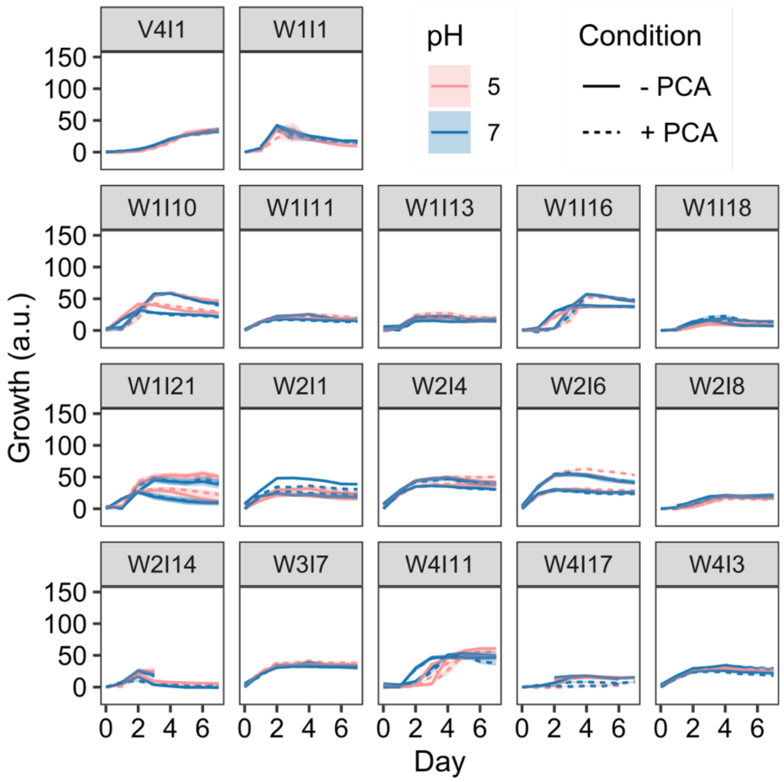


**Figure S7: Growth of Proteobacteria strains over time with and without PCA.**
Growth was quantified as described in the Methods. Solid lines represent the growth of spotted cultures on PCA-free agar, and dashed lines represent the growth of spotted cultures on agar containing 100 µM PCA. Blue represents growth at pH 7.3 and pink represents growth at pH 5.1. Data points are the mean of three technical replicates from a representative biological replicate for each strain, and the shaded ribbon represents the standard deviation. Different lines of the same color and line type (dashed vs. solid) represent separate biological replicates. For some biological replicates, data were collected only for a subset of days.

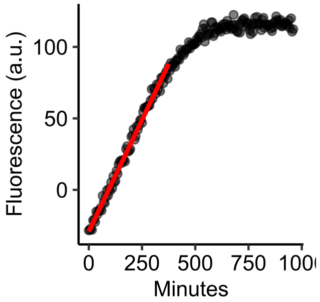


**Figure S8: Derivation of PCA reduction rate from fluorescence readings over time.**
This figure shows an example plot of fluorescence readings for reduced PCA over time for a single strain. Linear regression was performed to find the line of best fit to the range of the data that was judged by eye to be most linear, as depicted by the red line.

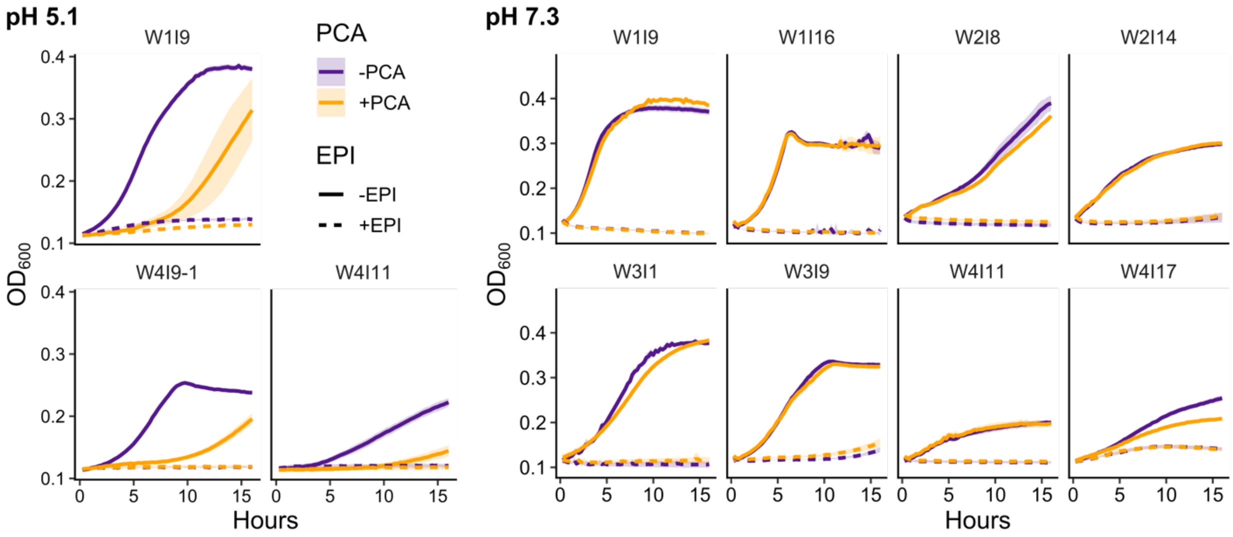


**Figure S9: Strain-specific synergistic toxicity of efflux pump inhibitors.**
Growth curves of strains for which treatment with a combination of the efflux pump inhibitors (EPI) reserpine and PAβN was toxic independently of PCA treatment. In all panels, data points are the mean of three biological replicates and the shaded ribbon represents the standard deviation. The concentrations of reserpine and PAβN used for each strain can be found in Table S2.

**Table S1: Strains used in this study.**

This table contains two tabs: **A.** Strains isolated in this study (wheat field isolates). **B.** Strains acquired from publicly available culture collections. For strains isolated in this study, an asterisk next to the percent identity to the closest BLAST hit indicates that the match was on the basis of only the reverse sequencing read of the 16S rRNA gene, due to multiple products in the forward read. Strains that are not identified to species level had multiple equal top hits to different species in the NCBI 16S rRNA gene sequence database.

**Table S2: Strain-specific concentrations of efflux pump inhibitors.**

This table lists the concentrations of reserpine and PAβN used for each strain in the experiments testing how efflux pump inhibitors (EPIs) affect susceptibility to PCA. The concentrations were optimized by determining the highest concentration of each EPI that, when administered separately, did not affect growth in the absence of PCA.
